## Supplemental Materials for "EEG microstate transition cost correlates with task demands"


**Fig. S1. Selection of the best number of microstates using the cross-validation criterion.** (*left*) Fraction of total variance (GEV) explained by the microstates. (*center*) Residual noise. (*right*) Cross-validation (CV) as a function of the number of microstates (N states).

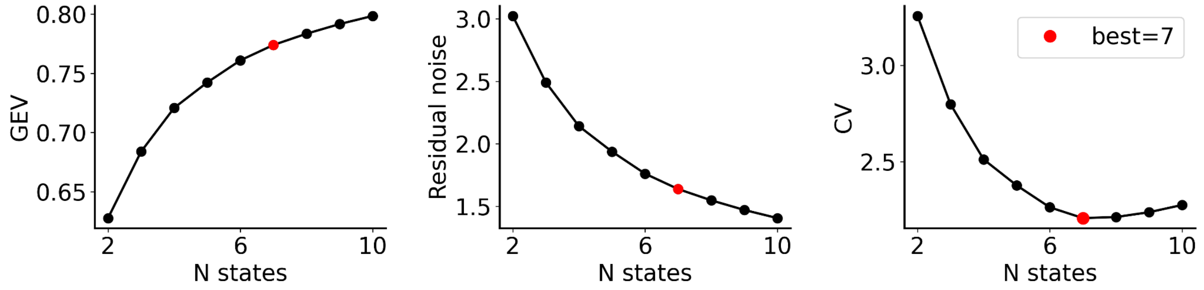


**Fig. S2. Behavioural results of the spatial Stroop task.**
Distribution of reaction times (RT) for the 44 participants during each task condition.

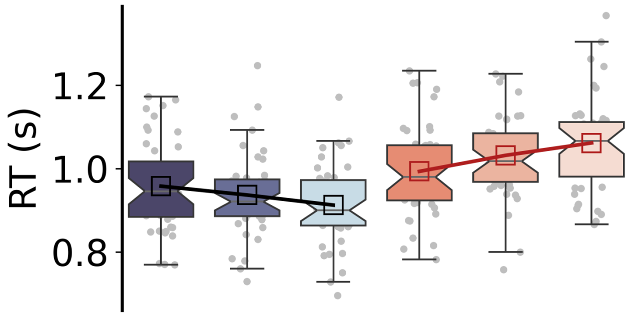


**Fig. S3. Global modulation of microstate distributions during the spatial Stroop task.**
Distribution of Kullback-Leibler divergence ($D_{KL}$) between the task ($\pi_{task}$) and resting ($\pi_{rest}$) for the 44 participants.

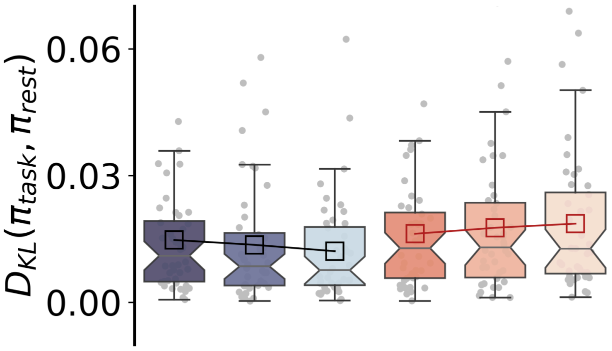

**Table S1. LMM results for microstate occurrences.**

For each microstate, a linear mixed model was implemented to test whether the change in the probability distribution during task execution compared to the resting state is modulated by the stimulus congruency, the PC level, and their interaction.

| State | Effect | *b* | *SE* | *t* | *df* | *p* | *d* | |
|---|---|---|---|---|---|---|---|---|
| A | Intercept | -0.018 | 0.004 | -4.92 | 44.00 | 0.0000 | -0.74 | * |
| A | PC | 0.000 | 0.001 | 0.15 | 62.03 | 0.8807 | 0.02 | |
| A | Congruency | 0.003 | 0.001 | 3.52 | 58.64 | 0.0008 | 0.46 | * |
| A | Congruency:PC | 0.005 | 0.002 | 2.51 | 101.65 | 0.0135 | 0.25 | * |
| B | Intercept | -0.006 | 0.003 | -2.12 | 44.00 | 0.0394 | -0.32 | |
| B | PC | -0.002 | 0.001 | -1.68 | 49.60 | 0.0985 | -0.24 | |
| B | Congruency | 0.005 | 0.001 | 5.14 | 48.49 | 0.0000 | 0.74 | * |
| B | Congruency:PC | 0.005 | 0.002 | 2.47 | 101.01 | 0.0153 | 0.25 | * |
| C | Intercept | 0.013 | 0.003 | 4.16 | 44.00 | 0.0001 | 0.63 | * |
| C | PC | 0.002 | 0.001 | 1.37 | 47.80 | 0.1770 | 0.20 | |
| C | Congruency | -0.005 | 0.002 | -3.07 | 44.56 | 0.0036 | -0.46 | * |
| C | Congruency:PC | -0.004 | 0.002 | -1.63 | 105.23 | 0.1065 | -0.16 | |
| D | Intercept | 0.010 | 0.003 | 3.61 | 44.00 | 0.0008 | 0.54 | * |
| D | PC | 0.000 | 0.001 | -0.10 | 98.38 | 0.9219 | -0.01 | |
| D | Congruency | -0.002 | 0.001 | -1.58 | 48.74 | 0.1196 | -0.23 | |
| D | Congruency:PC | -0.004 | 0.002 | -1.64 | 67.94 | 0.1046 | -0.20 | |
| E | Intercept | -0.012 | 0.003 | -3.96 | 44.00 | 0.0003 | -0.60 | * |
| E | PC | 0.000 | 0.001 | -0.15 | 44.11 | 0.8841 | -0.02 | |
| E | Congruency | 0.003 | 0.001 | 3.13 | 44.19 | 0.0031 | 0.47 | * |
| E | Congruency:PC | 0.004 | 0.002 | 2.47 | 101.53 | 0.0151 | 0.25 | * |
| F | Intercept | -0.007 | 0.004 | -1.81 | 44.00 | 0.0767 | -0.27 | |
| F | PC | 0.000 | 0.002 | -0.24 | 44.12 | 0.8144 | -0.04 | |
| F | Congruency | 0.005 | 0.001 | 4.65 | 44.16 | 0.0000 | 0.70 | * |
| F | Congruency:PC | 0.001 | 0.002 | 0.24 | 90.99 | 0.8140 | 0.02 | |
| G | Intercept | 0.021 | 0.002 | 10.09 | 44.00 | 0.0000 | 1.52 | * |
| G | PC | 0.001 | 0.002 | 0.39 | 44.25 | 0.6957 | 0.06 | |
| G | Congruency | -0.010 | 0.001 | -7.11 | 44.24 | 0.0000 | -1.07 | * |
| G | Congruency:PC | -0.006 | 0.003 | -2.50 | 105.04 | 0.0141 | -0.24 | * |

*Notes: b*, estimated coefficient; *SE*, standard error; *df*, degrees of freedom (estimated using the Satterthwaite method); The asterisks indicate the significant results after correction for multiple tests, applied using the false discovery rate approach.

### Text S1. Modified k-means clustering.

The modified k-means algorithm, introduced in [Pascual-Marqui et al., 1995], involves the traditional k-means clustering approach. However, it incorporates polarity-invariant topographical maps of the prototype microstates and considers the activation levels of these microstates, specifically their strength at each time point. In mathematical terms, each instantaneous activity map $x_t$ is assigned to the microstate index that minimizes the orthogonal squared Euclidean distance, $k_t = argmin_k\, d^2_{kt}$, where $d^2_{kt} = x_t^T x_t - (x_t^T a_k)^2$. It turns out to be equivalent to maximizing the spatial correlation. To define the best number of clusters, we employ the cross-validation criterion, which minimizes $CV = \sigma^2 (\frac{n_{ch}-1}{n_{ch}-K-1})^2$, where $\sigma^2 = \frac{\sum_t x_t^T x_t - (x_t^T a_t)^2}{T(n_{ch}-1)}$ is an estimator of the variance of the residual noise.
